# Longitudinal Changes in Auditory and Reward Systems Following Receptive Music-Based Intervention in Older Adults

**DOI:** 10.1101/2021.07.02.450867

**Authors:** Milena Aiello Quinci, Alexander Belden, Valerie Goutama, Dayang Gong, Suzanne Hanser, Nancy J. Donovan, Maiya Geddes, Psyche Loui

## Abstract

Listening to pleasurable music is known to engage the brain’s reward system. This has motivated many cognitive-behavioral interventions for healthy aging, but little is known about the effects of music-based intervention (MBI) on activity and connectivity of the brain’s auditory and reward systems. Here we show preliminary evidence that brain network connectivity can change after receptive MBI in cognitively unimpaired older adults. Using a combination of whole-brain regression, seed-based connectivity analysis, and representational similarity analysis (RSA), we examined fMRI responses during music listening in older adults before and after an eight-week personalized MBI. Participants rated self-selected and researcher-selected musical excerpts on liking and familiarity. Parametric effects of liking, familiarity, and selection showed simultaneous activation in auditory, reward, and default mode network (DMN) areas. Functional connectivity within and between auditory and reward networks was modulated by participant liking and familiarity ratings. RSA showed significant representations of selection and novelty at both time-points, and an increase in striatal representation of musical stimuli following intervention. An exploratory seed-based connectivity analysis comparing pre- and post-intervention showed significant increase in functional connectivity between auditory regions and medial prefrontal cortex (mPFC). Taken together, results show how regular music listening can provide an auditory channel towards the mPFC, thus offering a potential neural mechanism for MBI supporting healthy aging.

## 1 Introduction

Recent interest in music has burgeoned as a tool for restoring function in the aging brain [1]. The possibility that music can maintain or even restore cognitive and/or emotional function in old age hinges upon the observation that when listening to pleasurable music, areas in the auditory network are functionally connected with areas in the dopaminergic reward system, specifically medial prefrontal cortex (mPFC) as well as dorsal and ventral striatum [2-5]. The reward system is involved in motivated behavior [6] and is active during the processing of biologically relevant stimuli such as food, as well as cues that are tightly linked to biological stimuli, such as money [7]. Reward system areas are also involved in the learning of associations between stimuli and their subjective value [3, 8-10]. This activity and connectivity of the reward system declines in old age, as supported by evidence for age-related decline in pre- and post-synaptic dopamine function in human striatum and midbrain [11-13]. On the other hand, older adults have preserved brain activation of the striatum (caudate) to positively-valenced stimuli compared to younger adults [14]. Since music can be a rich source of positively-valenced stimuli, using music as a means by which to stimulate the reward system may have enduring effects towards old age.

Despite their known coactivity during music listening, little is understood about how connectivity between auditory and reward areas are modulated by familiarity and self-selection. Understanding the stimulus selection parameters that influence auditory-reward connectivity will inform studies on music, health, and well-being, with especially widespread implications for music therapy. As the evolutionary bases and social roles of music are subserved neurobiologically by the ability of music to stimulate the reward system [15], studying auditory-reward connectivity during music listening may inform these evolutionary functions of music as well.

In addition to activating the reward system, the experience of music engages multiple other brain systems such as sensorimotor and executive control networks [16, 17], and is fundamentally psychologically intertwined with the listener’s knowledge, autobiographical memories, and understanding of the intentions of the composer and the performer, through its structure and performance [18-20]. These rich experiences are supported by activity in the default mode network (DMN), which encompasses the mPFC (which is also part of the reward network), along with the posterior cingulate cortex (PCC) and temporoparietal junction (TPJ) or angular gyrus. The DMN is involved during self-generated thought [21] and is disrupted in activity and connectivity in multiple neuropsychiatric disorders and conditions such as depression and learned helplessness [22].

Agency in the selection of music, or the ability to choose the music that one listens to, may provide the listener with a psychological platform to link the consequences of one’s musical choices to perceptual predictive processes [23], giving rise to a relatively active mode of listening [24] that has implications for the health-relevance of music and the use of music in therapy [25]. Listening to self-selected music reduces anxiety and improves task performance and enjoyment, compared to music selected by an experimenter [26]. While previous fMRI studies suggest that the reward network and DMN are likely involved [27], there is little direct evidence for the involvement of these networks in self-selection of music listening per se. Since the mPFC is both within the DMN and in the reward network, testing how self-selected music activates these disparate networks may inform neuroscientific understanding about how the reward and default networks interact in the brain.

Here we present the first report of task fMRI results on music listening before and after an ongoing music-based intervention (MBI) in cognitively unimpaired, community-dwelling older adults. MBIs are therapeutic strategies facilitated by music, and can be classified as either receptive if they are primarily focused on music listening, or interactive if they involve active engagement in music making activities [28]. Receptive MBIs have been shown to reduce depressive symptoms in individuals with clinical depression [29, 30] and to improve quality of life and emotional well-being, while decreasing anxiety for people with dementia [31]. Relatedly, an eight-week intervention on mindfulness-based stress reduction has shown changes in reward-related fMRI activity and connectivity [32]. Thus, we developed an eight-week receptive MBI program, based on previous research with older adults [33], and collected neuropsychological and neuroimaging measures both prior to and after participation in the intervention. Here we report preliminary evidence from the fMRI portion of the larger study. Through this project, we aim to investigate the effects of music liking, familiarity, and self-selection on reward processing before and after intervention, through a combination of whole-brain regression, seed-based analysis, and representational similarity analysis (RSA).

Our predictions (preregistered here:https://osf.io/qsevr/) are twofold: first, we predict that brain responses to musical stimuli will be sensitive to differences in liking, familiarity, and self-selection. Specifically, from a whole-brain analysis, we expect higher brain activity when listening to musical stimuli that are well-liked, familiar, and self-selected. Second, we predict that the activity and connectivity within and between auditory and reward areas, as observed in region-of-interest (ROI) analyses, will be sensitive to differences in liking, familiarity, and self-selection. Finally, as an additional exploratory hypothesis, we expect some experience-dependent changes in how the brain responds to music over time, to be observed by comparing activity and connectivity in auditory and reward regions before and after MBI.

## 2 Materials and Methods

### 2.1 Participants

Twenty-two older adults met inclusion criteria to participate. Three of them dropped out during the first session for various reasons related to the MRI portion of the study, and three were lost-to-follow-up after the second session as a result of the COVID-19 pandemic, resulting in a final sample of 16 older adults (8 males and 8 females) with the mean age of 66.38 (*SD*=8.74). Recruitment took place through craigslist.org and through coauthors (MG and ND) at Brigham and Women’s Hospital. Participants were included if they 1) were at least 50 years old, 2) passed MRI screening, and 3) had had no more than mild hearing loss as defined by a pure-tone audiogram showing less than 40 dB hearing loss, in accordance with the American Speech-Language-Hearing Association. Participants were excluded if they 1) changed medications within 6 weeks of screening; 2) had a history of psychotic or schizophrenic episodes, major neurological diagnosis, or other medical condition that might impair cognition; 3) had a history of chemotherapy within the past 10 years; or 4) experienced serious physical trauma or were diagnosed with a serious chronic health condition requiring medical treatment and monitoring within 3 months of screening. Participants reported between 0-10+ years of musical experience as assessed using the Gold-MSI (Mullensiefen et al, 2014). Eight participants reported receiving 0 years of musical training, one reported 1 year, three reported 3-5 years, one reported 6-9 years, and two participants reported 10+ years. Fifteen of the participants listed English as their first language and one listed Russian. Participants were also asked if they were fluent in any other languages, to which one participant indicated fluency in Spanish. Recruitment for this study was significantly affected the COVID-19 pandemic as many individuals were nervous about participating in in-person research (see section 2.3 for further details). Participants were compensated for their time. This study was approved by the Northeastern University Institutional Review Board. Informed consent was obtained from all participants. All research was performed in accordance with relevant guidelines/regulations, and in accordance with the Declaration of Helsinki.

### 2.2 Procedure

Prior to their enrollment, participants completed a pre-screening by telephone. Participants were screened for the inclusion and exclusion criteria and completed the Telephone Interview for Cognitive Status (TICS) [34] with a score of ≥ 31/41 considered cognitively normal and an MRI pre-entry screening form. They were also asked to provide the researcher with the names of six musical pieces they enjoy listening to. Participants who passed pre-screening were invited to the lab for a pre-intervention session, which included a battery of neuropsychological tests and behavioral measures, an MRI, and a blood draw. The neuropsychological tests were administered by a trained researcher, and the behavioral measures were completed through an online survey. Later that week, participants were invited back to the lab to meet with a music therapist for an assessment of their musical preferences and creation of two personalized playlists for the MBI (see section 2.4). At the end of the 8-week intervention period (12 weeks for the 2 participants enrolled during COVID-19; see Section 2.3), participants returned to the lab for a post-MBI session to complete the same battery of neuropsychological tests and behavioral measures administered during the first session, as well as a post-intervention MRI and blood draw. See Figure 1 for a timeline of the procedure. For the present manuscript, we report data from the task fMRI portion of the study; data from the neuropsychological battery and the behavioral measures will be reported separately.

**Figure 1.**
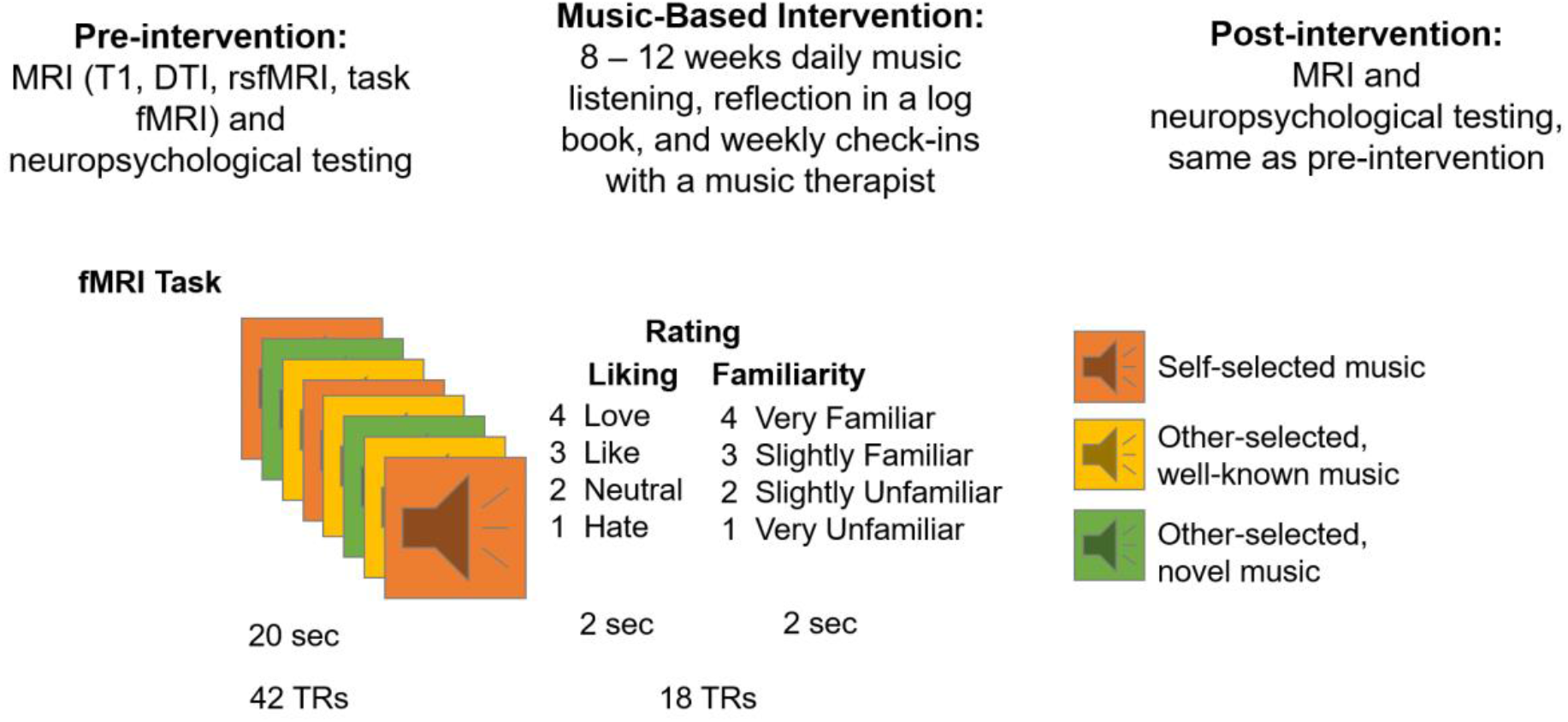
Study timeline and fMRI experiment design.

### 2.3 Special Considerations due to COVID-19

Data collection for this study began in July of 2019, and had been completed per protocol on 10 participants before the United States entered a COVID-19-related lockdown. Two participants included in the final sample were actively undergoing MBI when the lockdown began in March 2020. For those two participants, the music intervention was extended to 12 weeks rather than the typical 8 weeks, to complete the intervention as intended. Resumption of Research Plan was developed and approved in consultation with the Institutional Review Board (IRB) of Northeastern University. Removing those two participants did not alter the overall pattern of results, thus they were analyzed along with the rest of the sample. The remaining 4 participants in the final sample were enrolled during the COVID-19 pandemic, and underwent an 8-week MBI and performed both pre- and post-intervention testing in compliance with the COVID-specific Resumption of Research plan.

For our COVID-specific Resumption of Research plan: consenting and neuropsychological testing were done via a HIPAA-compliant version of Zoom to minimize the amount of time participants spent in the lab for in-person data collection. Participants were called one day before their scheduled session, screened using a COVID-19 pre-screener based on the COVID-19 CDC guidelines, and were provided with the lab’s IRB-approved COVID-19-compliant guidelines. If participants answered no to all of COVID-19 pre-screener questions, they were invited to come in for their session the next day. After arriving for their session, participants were screened again using the same COVID-19 pre-screener and their temperatures were taken. If participants answered no to all of the questions and had a temperature reading of less than 100.4° F, then they were invited into the lab. Participants and researchers wore full personal protective equipment (PPE: surgical masks, face shields, lab coats, and surgical gloves) and were socially distanced as much as possible during the onsite portions of the study, and the lab was disinfected before and after each session. In the MRI, when alone in the scanner, participants were allowed the option of taking their masks off. Recent fMRI studies suggest that wearing a mask during the scan does not significantly affect task activation [35].

### 2.4 Music-Based Intervention

The current approach to MBI was based on a previously developed music therapy protocol that was shown by a randomized, controlled trial to be successful in improving symptoms of depression, anxiety, and distress in older adults diagnosed with mild or moderate depression [33]. Subsequent research applied this intervention to other clinical settings, and condensed the music used into two manageable playlists, composed of energizing and relaxing music [36-38]. Thus, MBI was personalized in that each participant met (in-person or virtually) with a board-certified music therapist to help curate playlists. Participants built and modified their playlists for the MBI independent of what pieces they initially selected to listen to in the scanner. Prior to the start of the MBI, a Premium YouTube Music account was created for each participant for music listening. At the start of the MBI, participants met with a board-certified music therapist who familiarized them with the YouTube Music platform and assessed their music preferences. An energizing playlist and a relaxing playlist, of approximately 25 pieces of music each, were compiled in consultation with the music therapist, beginning with the six favorite pieces selected during pre-screening. Participants built and modified their playlists for the MBI independent of what pieces they initially selected to listen to in the scanner. Some of the chosen playlists used in MBI (the energizing and relaxing playlists) overlapped with the 6 songs used in the scanner, but this was not a requirement for the MBI or for the self-selection of songs to be included in the scanner task. On average, participants’ energizing playlist contained 2.25 (std 2.21) of their self-selected pieces at the beginning of the intervention, and 1.69 (1.54) at the end of intervention. Participants’ relaxing playlist contained 2.44 (2.22) of their self-selected pieces at the beginning of the intervention, and 2.38 (2.09) at the end of intervention. The music therapist instructed participants to listen to their choice of selections from one or both of their playlists for one hour daily, over the course of the next 8 weeks. They were asked to pay attention to how the music affected their moods and emotions, while noting any memories, images, or associations with the music. After each focused listening experience, participants recorded these reflections and self-observations in a journal. Once a week, the music therapist called the participant to discuss their compliance and responses to the daily listening, and to help them add or remove pieces from their playlists. Prior to COVID-19, the start of the MBI occurred in person; however, during COVID-19, participants met with the music therapist over Zoom.

### 2.5 fMRI Task

The fMRI task consisted of 24 trials altogether. In each trial, participants were first presented with a musical stimulus (lasting 20 seconds), then they were given the task of rating how familiar they found the music to be (familiarity rating lasted 2 seconds), and how much they liked the music (liking rating also lasted 2 seconds). Musical stimuli for the MRI task consisted of 24 different audio excerpts. Each auditory stimulus was from one of the following three categories: participant self-selected music (6/24 stimuli), other-selected (researcher-selected) music including well-known excerpts spanning multiple musical genres [39](10/24 stimuli) and novel music spanning multiple genres (8/24 stimuli). A list of the researcher-selected musical selections is given in Supplementary Materials Table S1. Stimuli were presented in a randomized order, and participants made ratings of familiarity and liking on the scales of 1 to 4: for familiarity: 1=very unfamiliar, 2=unfamiliar, 3=familiar, 4=very familiar; for liking: 1=hate, 2=neutral, 3=like, 4=love. Participants made these ratings by pressing a corresponding button on a button-box (Cambridge Research Systems) inside the scanner. Participants wore MR-compatible over-the-ear headphones (Cambridge Research Systems) over musician-grade silicone ear plugs during MRI data acquisition. The spatial mapping between buttons and the numerical categories of ratings were counterbalanced between participants to reduce any systematic association between familiarity or liking and the motor activity resulting from making responses. This fMRI task was completed before and after the MBI.

### 2.6 fMRI Data Acquisition

Images were acquired using a Siemens Magnetom 3T MR scanner with a 64-channel head coil at Northeastern University. For task fMRI data, continuous acquisition was used for 1440 volumes with a fast TR of 475 ms, for a total acquisition time of 11.4 minutes. Forty-eight axial slices (slice thickness = 3 mm, anterior to posterior, z volume = 14.4 mm) were acquired as echo-planar imaging (EPI) functional volumes covering the whole brain (TR = 475 ms, TE = 30 ms, flip angle = 60°, FOV = 240mm, voxel size = 3 × 3 × 3 mm^3^).

T1 images were also acquired using a MPRAGE sequence, with one T1 image acquired every 2400 ms, for a total task time of approximately 7 minutes. Sagittal slices (0.8 mm thick, anterior to posterior) were acquired covering the whole brain (TR = 2400 ms, TE = 2.55 ms, flip angle = 8°, FOV= 256, voxel size = 0.8 × 0.8 × 0.8 mm^3^). Resting state and diffusion tensor imaging data were also collected, but as they are outside the scope of the present study, they will be reported in another manuscript.

### 2.7 Data Analysis

#### 2.7.1 Preprocessing

Task fMRI data was preprocessed using the Statistical Parametric Mapping 12 (SPM12) software [40] with the CONN Toolbox [41]. Preprocessing steps included functional realignment and unwarp, functional centering, functional slice time correction, functional outlier detection using the artifact detection tool, functional direct segmentation and normalization to MNI template, structural centering, structural segmentation and normalization to MNI template, and functional smoothing to an 8mm gaussian kernel [42]. Denoising steps for fMRI data included white matter and cerebrospinal fluid confound correction [43], and bandpass filtering to 0.008– 0.09 Hz.

#### 2.7.2 Univariate Whole-Brain Analysis

Response data from the fMRI task were imported into R Studio, and onset values were extracted for each trial, including its liking rating (hate, neutral, like, love), familiarity rating (very unfamiliar, unfamiliar, familiar, very unfamiliar), and music condition (self-selected, other-selected well-known, other-selected novel). Pre-intervention, four participants did not have onsets for the familiar condition (four participants did not feel that any of the stimuli were “familiar” to them), and four participants did not have onsets for the hate condition. Post-intervention, two participants were missing onsets for unfamiliar, one for familiar, one for neutral, and one for hate. Additionally, one of our participants post-intervention was missing all data for familiarity ratings due to a technical error.

First- and second-level analyses were completed in SPM12 and visualized through CONN. For each participant, data were converted from 4D to 3D images, resulting in 1440 scans. The model was specified using the following criteria: interscan interval = 0.475 seconds, microtime resolution = 16, microtime onset = 8, duration = 42. Only data from the time while the participant was listening to the musical excerpt (rather than when the participant was making the familiarity and liking ratings) were included in this model. At the first-level, the following contrasts were extracted both pre- and post-intervention: the parametric effect of liking (linear contrast with all liking ratings), the parametric effect of familiarity (linear contrast with all familiarity ratings), and self-selected > other-selected music. The resulting first-level contrasts were then analyzed using a one-sample t-test across all participants at the second level. Results from the second-level analyses were statistically corrected using a cluster and voxel threshold of *p* <0.05 (FDR-corrected). Univariate pre-intervention and post-intervention effects of liking, familiarity, and self-selection did not differ at the whole-brain FDR-corrected level. We thus combined pre- and post-intervention scans in a conjunction analysis to show the effects that were common to both stages of the longitudinal study [44]. Whole-brain results were rendered to a standard MNI brain.

#### 2.7.3 Seed-Based Connectivity Analysis

Seed-based connectivity analysis was performed using the CONN toolbox to determine changes in functional connectivity after participation in MBI. Seed ROIs consisted of Auditory and Reward networks defined by previous work in our lab [45, 46], with the effect of session used as a between-conditions contrast (post-intervention > pre-intervention). Significant clusters from these comparisons were then processed as new ROI masks, and beta time-series for these ROIs were extracted and compared trial-by-trial across the four liking and familiarity ratings for each subject.

#### 2.7.4 Auditory and Reward Network Connectivity

Beta-weights for a series of auditory and reward network ROIs [47] were extracted from participant’s first-level SPM.mat files using the Marsbar Toolbox [48]. Values associated with each liking rating (hate, neutral, like, love) and each familiarity rating (very unfamiliar, unfamiliar, familiar, very familiar) were separated for each participant and averaged across all stimuli with that particular rating. For each rating, beta-weights were correlated across all auditory regions (auditory-auditory), across all reward regions (reward-reward), and between auditory and reward regions (auditory-reward).

#### 2.7.5 Representational Similarity Analysis

Representational dissimilarity matrix (RDM) definition and comparisons were performed in MATLAB (Mathworks, R2020b) using tools from the RSA toolbox [49]. RDMs used in this study were a combination of fMRI-derived RDMs and model RDMs. Selection and Novelty RDMs were generated by assigning each stimulus either a 1 or a 0 depending on if it was one of the six participant-selected stimuli or one of the 16 well-known stimuli (including self-selected and researcher-selected excerpts), respectively, and then subtracting each pair of stimulus values to form a binary difference matrix. The model RDM of Familiarity was defined as the difference in participant familiarity ratings between each pair of stimuli, and similarly the model of Liking used subject liking ratings.

fMRI-derived RDMs were generated using ROI-specific beta timeseries information for each stimulus. Each fMRI-derived RDM was defined as one minus the correlation coefficient of each pair of stimuli’s average beta timeseries for a given ROI. Regions of interest included in this analysis consist of the 14 networks defined by the Stanford FIND lab atlas [50], as well as 12 hypothesis-driven ROIs, including ROIs for bilateral anterior and posterior superior temporal gyrus (STG), bilateral Heschl’s Gyri (HG), Nucleus Accumbens (NAcc), and supplementary motor area (SMA), the mPFC, and the PCC derived from CONN Default and Networks Atlas. Finally, we included auditory and reward networks as defined in a previous study [45].

First-level, subject specific RDMs were generated for each ROI at both pre-intervention and post-intervention timepoints, and then compared to model RDMs. ROI to model comparisons were achieved through two independent analysis pipelines. In the first analysis pipeline, each first-level fMRI-derived RDM was compared to models of Selection, Novelty, Liking, and Familiarity through 10,000-fold bootstrap resampling [49, 51] to generate relatedness values for each subject between each ROI-Model pair. These values then underwent 50,000 fold permutation T-max testing in order to determine significant ROI Model relationships at each session, and to determine any significant differences between sessions. The second analysis pipeline averaged first-level RDMs in order to form second-level, subject averaged RDMs. These second-level RDMs were then visualized using second order multidimensional scaling [49].

## 3 Results

### 3.1 Behavioral Results

Table 1 shows the means and SEs of liking and familiarity ratings of different stimulus types, before and after the intervention. We ran two repeated-measures ANOVAs, one for the liking ratings and one for the familiarity ratings, to determine the main effect of time (pre-intervention vs post-intervention) rating and the time x rating interaction.

**Table 1.** Descriptive Statistics for Liking and Familiarity Behavioral Data

| **Liking** |  |  |
| --- | --- | --- |
| Condition | Mean | SE |
| Time |  |  |
| Pre | 2.85 | 0.103 |
| Post | 2.70 | 0.080 |
| Stimulus Type |  |  |
| Self-Selected | 3.77 | 0.079 |
| Well-Known | 2.74 | 0.120 |
| Novel | 1.81 | 0.138 |
| Time x Stimulus Type |  |  |
| Pre Self-Selected | 3.69 | 0.139 |
| Pre Well-Known | 2.88 | 0.128 |
| Pre Novel | 1.98 | 0.153 |
| Post Self-Selected | 3.85 | 0.059 |
| Post Well-Known | 2.60 | 0.126 |
| Post Novel | 1.64 | 0.141 |
| **Familiarity** |  |  |
| Condition | Mean | SE |
| Time |  |  |
| Pre | 2.42 | 0.086 |
| Post | 2.45 | 0.057 |
| Stimulus Type |  |  |
| Self-Selected | 3.82 | 0.076 |
| Well-Known | 2.24 | 0.115 |
| Novel | 1.23 | 0.074 |
| Time x Stimulus Type |  |  |
| Pre Self-Selected | 3.70 | 0.141 |
| Pre Well-Known | 2.26 | 0.133 |
| Pre Novel | 1.30 | 0.097 |
| Post Self-Selected | 3.94 | 0.045 |
| Post Well-Known | 2.23 | 0.118 |
| Post Novel | 1.18 | 0.070 |

#### 3.1.1 Liking

For liking ratings, there was a significant main effect of stimulus type (F(2, 30) = 103.8, p <0.001, 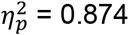). Average ratings for self-selected stimuli were significantly higher than for well-known and novel stimuli (p <0.001 Bonferroni-corrected), and ratings for the well-known stimuli were higher than for novel stimuli (p <0.001 Bonferroni-corrected). There was no significant main effect of time (F(1, 15) = 4.330, p = 0.055, 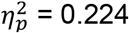). There was, however, a significant time x stimulus type interaction (F(1.4, 30) = 7.18, p = 0.009, 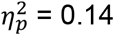), where well-known and novel stimuli were rated higher pre-intervention than post-intervention, but ratings for self-selected stimuli were rated higher post-than pre-intervention.

#### 3.1.2 Familiarity

For familiarity ratings, there was a significant main effect of stimulus type (F(2,28) = 307.7, p <0.001, 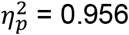). Bonferroni-corrected pairwise comparisons revealed that average familiarity ratings for self-selected stimuli were significantly higher than those for well-known (p <0.001) and novel stimuli (p <0.001), and familiarity ratings for the well-known stimuli were greater than for novel stimuli (p <0.001). There was no significant main effect of time (F(1, 14) = 0.343, p = 0.567, 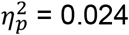), and no significant time x stimulus type interaction (F(1.4, 28) = 2.590, p = 0.114, 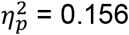).

### 3.2 Whole-Brain Univariate Effects of Liking, Familiarity, and Self-Selection

Our first hypothesis, i.e. that brain responses to musical stimuli would be sensitive to differences in liking, familiarity, and self-selection, was tested using parametric contrasts on liking and familiarity ratings, and a contrast between self-selected and other-selected music. Whole-brain effects for each liking rating, each familiarity rating, and each stimulus type can be found in https://neurovault.org/collections/TCPSFBMF/, and in Supplementary Materials Figure S1-S3.

#### 3.2.1 Parametric Effect of Liking

Figure 2A and Table 2 show the whole-brain corrected univariate, second-level parametric effects of liking ratings at the p <.05 voxelwise and clusterwise FDR-corrected level. A parametric effect of liking was observed in auditory regions including bilateral STG, superior temporal sulcus (STS), and middle temporal gyrus (MTG); and DMN regions (including mPFC, PCC, and TPJ/inferior parietal lobule (IPL)).

**Figure 2.**
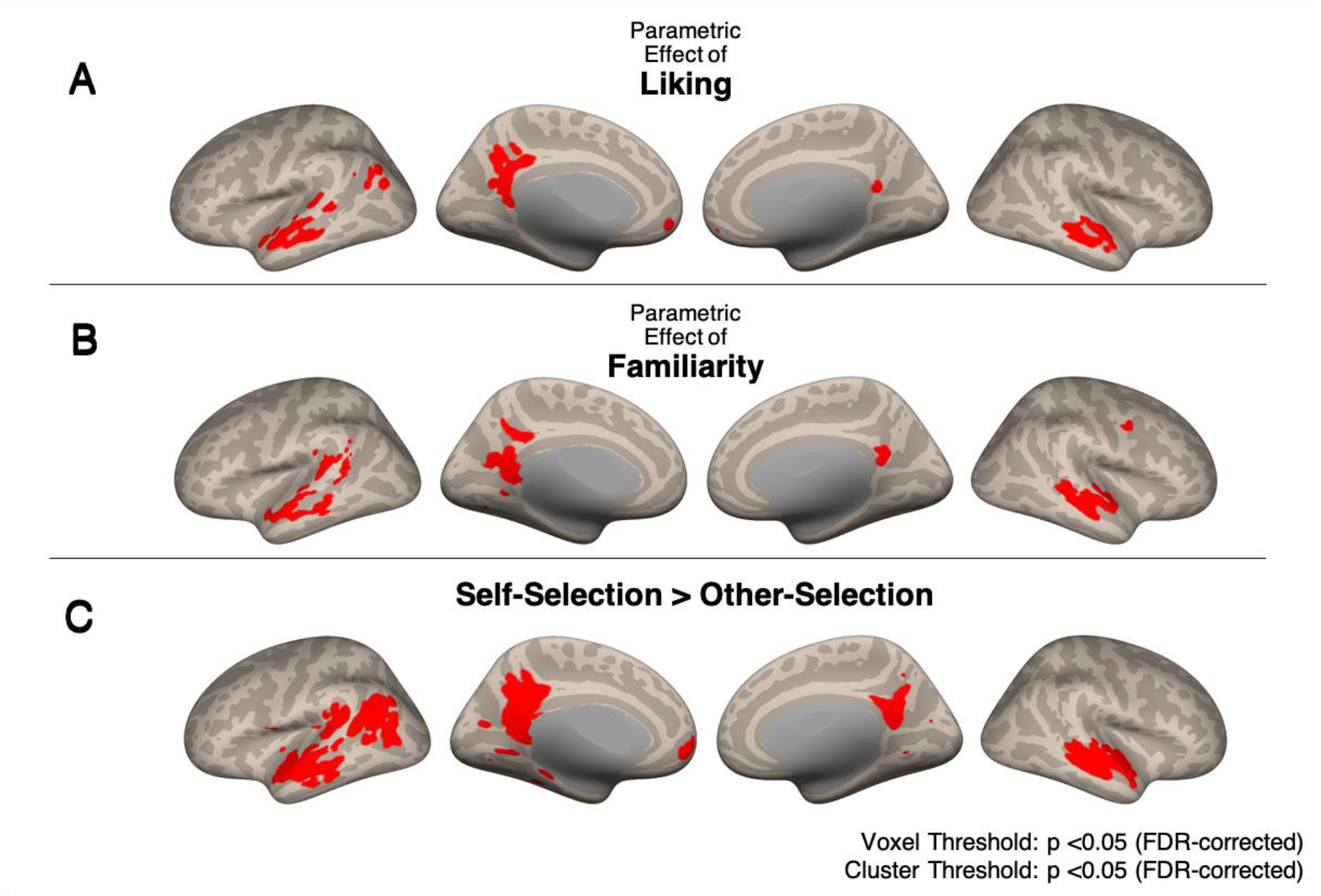
Conjunction analysis of pre- and post-intervention univariate whole-brain results. (A) Parametric effects of liking. Auditory (STG, STS, MTG), reward and DMN (mPFC and PCC) regions showed activity that varied parametrically with liking ratings. **(B) Parametric effects of familiarity**. Auditory and DMN regions showed activity that also varied parametrically with familiarity ratings. **(C) Self-selected > Other-selected music**. Greater widespread engagement was observed in auditory, reward, and DMN regions for self-selected music compared to other-selected music. All images are results of second-level analyses showing significant clusters at the p <.05 FDR-corrected level.

**Table 2.**
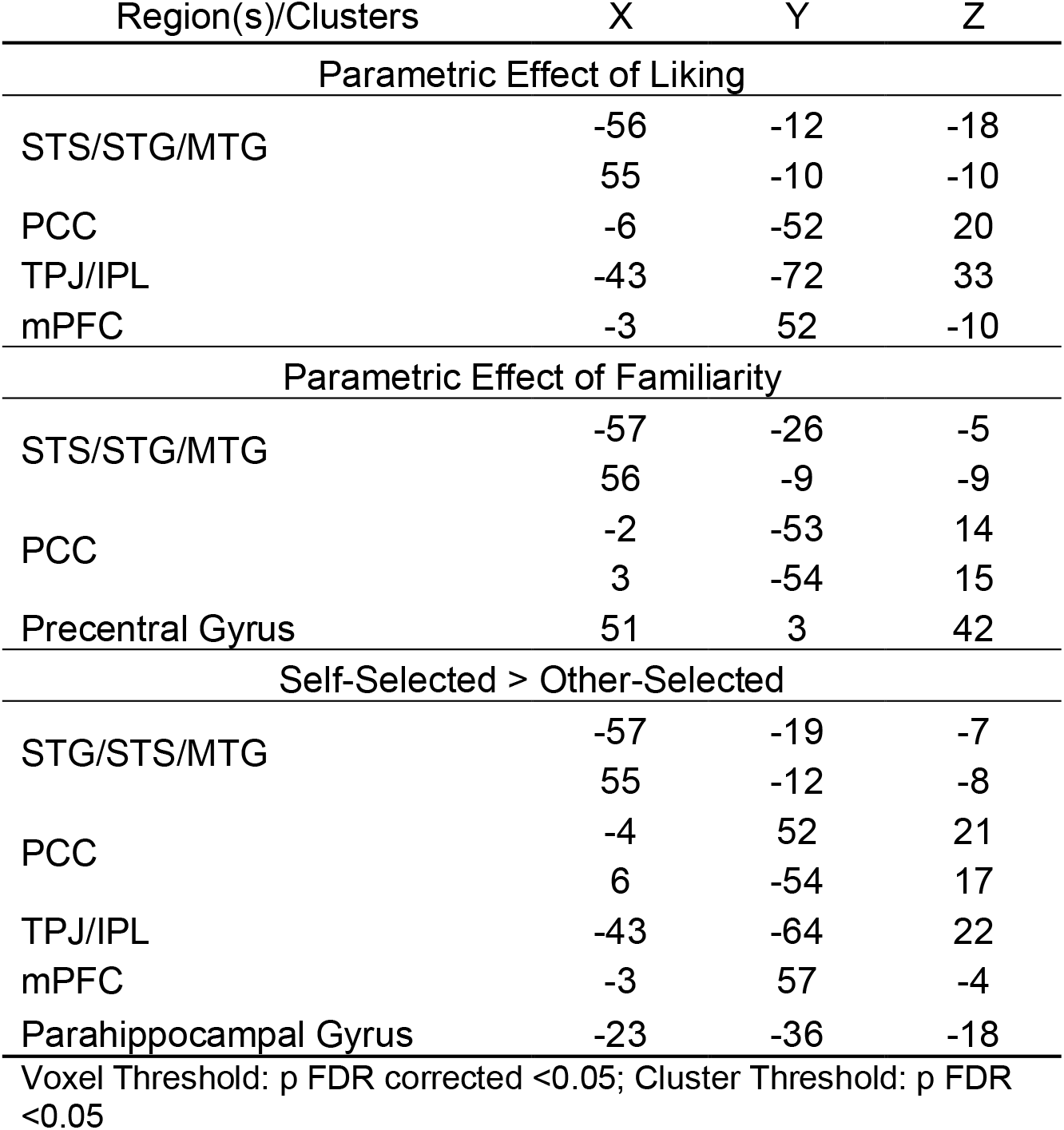
Conjunction Analysis of Pre- and Post-Intervention

#### 3.2.2 Parametric Effect of Familiarity

Figure 2B and Table 2 show the whole-brain corrected univariate, second-level parametric effects of familiarity ratings, again at the p <.05 voxelwise and clusterwise FDR-corrected level. A parametric effect of familiarity was observed in auditory regions (including bilateral STG, STS, and MTG); the PCC; and the precentral gyrus.

#### 3.2.3 Effect of Self vs. Other-Selection

Figure 2C and Table 2 show the whole-brain corrected univariate, second-level contrast of self-selected vs. other-selected music listening, again at the p <.05 voxelwise and clusterwise FDR-corrected level. Self-selected music, compared to other-selected music, elicited greater activation in the bilateral auditory areas (including STG, STS, and MTG); DMN (including mPFC, PCC, TPJ/IPL, and parahippocampal gyrus).

### 3.3 Seed-Based Connectivity: Post-intervention > Pre-intervention

Figure 3A shows pre- and post-intervention activity for the highest liking rating, showing significant activity in DMN areas post-intervention but not pre-intervention, suggesting some change in how the brain responds to music over time. To formally test our hypothesis of experience-dependent changes in activity and connectivity of auditory and reward regions, we conducted seed-based connectivity analyses, using auditory regions as the seed ROI and comparing its functional connectivity patterns before and after MBI. Figure 3B shows seed-based functional connectivity from the seed ROIs of the auditory network, including HG, STG, STS, and MTG (Wang et al., 2020). A between-sessions contrast (Post-intervention > Pre-intervention) on functional connectivity maps seeded from the auditory network showed a significant positive cluster in the mPFC at the p <.05 FDR-corrected level, indicating higher functional connectivity post-intervention than pre-intervention in this region. Thus, auditory areas increased in their connectivity to the mPFC after the intervention. Since this specific location within mPFC was the only region that increased its functional connectivity with the auditory network, we extracted the beta-series from this significant mPFC cluster for further ROI-based analyses.

**Figure 3.**
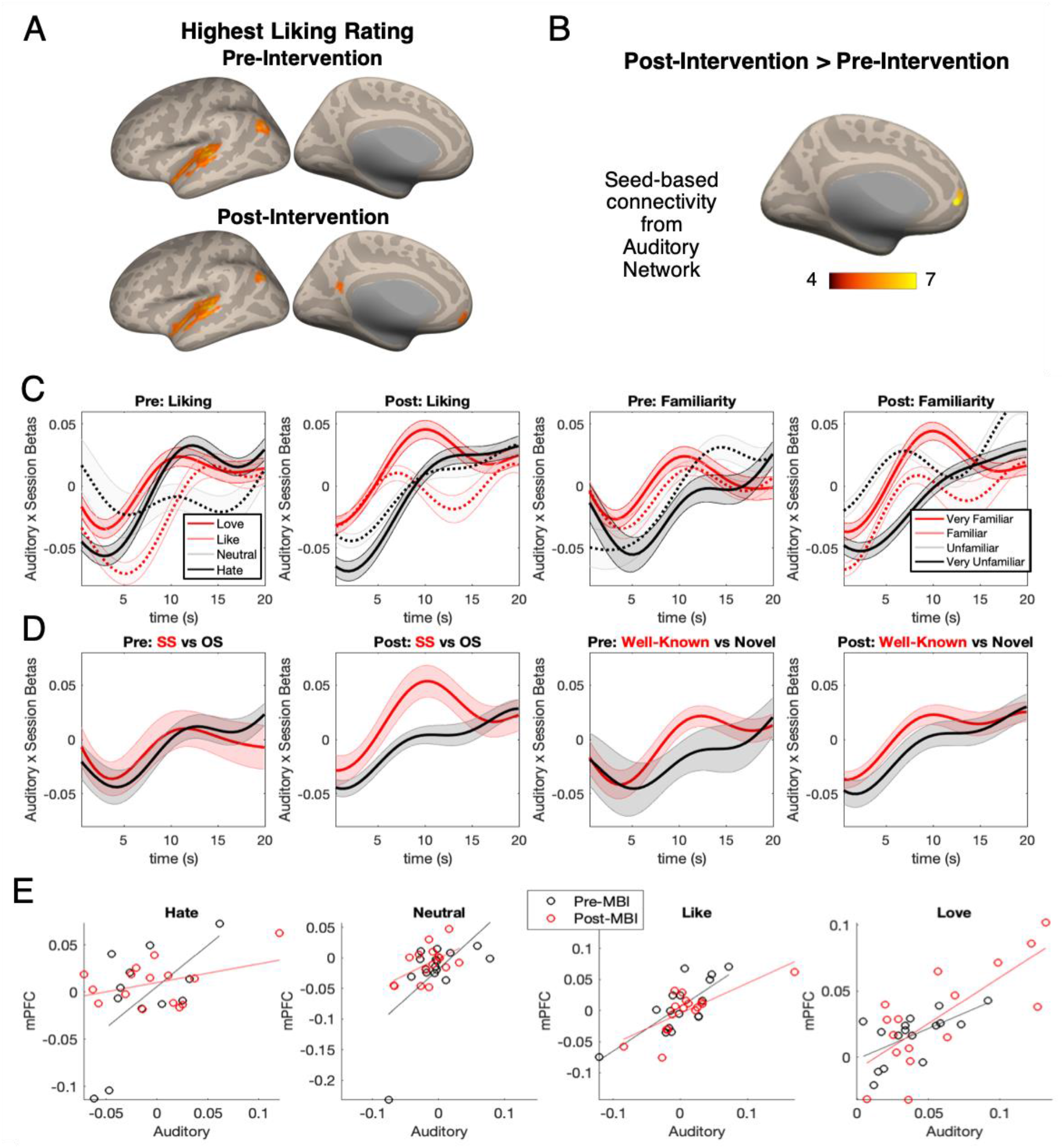
Seed-based connectivity from auditory ROIs. (A) Cortical regions active during musical stimuli that received the highest liking rating (“Love”) pre- and post-intervention, showing significant activity in mPFC post-intervention but not pre-intervention. Voxelwise and clusterwise FDR-corrected p<0.05. (B) The mPFC showed a significant effect of session (post-intervention > pre-intervention) in auditory network seed-based functional connectivity (voxel height p<0.001, uncorrected; cluster size p<0.05, FDR corrected). (C-D) Time-series data of ROIs extracted from the positive contrast in (B), representing post > pre-intervention increase in functional connectivity. (C) Beta series for different liking and familiarity ratings pre- and post-intervention, showing an increase in beta post-intervention, representing increased functional connectivity between auditory regions and mPFC, for loved and very familiar music. (D) Beta series for self-selected and other-selected, and for well-known and novel music listening, showing an increase in auditory-mPFC functional connectivity after intervention for self-selected music but not for other-selected music. This increase in functional connectivity was not observed when comparing well-known vs. novel music listening. (E) Correlations between auditory and mPFC activity in beta values for the four levels of liking ratings before and after intervention. Each data point is one subject. Comparing across the panels from “Hate” (lowest liking rating) to “Love” (highest liking rating) shows more correlated auditory and mPFC activity during high liking ratings compared to low liking ratings, with the between-subject correlation being highest after the intervention for stimuli that received higher ratings.

### 3.4 ROI analysis

To test our hypothesis that auditory and reward regions would be sensitive in their activity and connectivity to the self-selectedness of music, we extracted beta values from the auditory network [45] and the mPFC from the reward network. Figure 3E shows between-subject correlations between auditory and mPFC areas, pre- and post-intervention for the four levels of liking ratings. While auditory and mPFC activity were positively correlated across all conditions, the correlation was highest (r = 0.75) for musical stimuli that received a “Love” rating. Furthermore, the between-participants correlation between auditory and mPFC regions were higher post-intervention than pre-intervention, suggesting an increase in auditory-reward connectivity over time.

To address the functional connectivity of auditory and reward regions over the time-course of listening to music, we extracted the time-series of beta values (TR = .475 s) from the significant mPFC cluster identified in the auditory seed-based connectivity analysis above, and compared them across conditions and across sessions. While there was no clear difference between the conditions in pre-intervention beta-series, a clear pattern emerged between conditions in the post-intervention beta-series, such that at the midpoint of the 20-second-long musical stimuli, beta values were highest for self-selected, highly familiar, and well-liked stimuli (Figure 3CD). In other words, auditory-reward connectivity increased over time especially for well-liked, familiar, and self-selected musical excerpts, but did not increase over time for well-known vs. novel pieces of music.

### 3.5 Relating Behavioral Ratings to Auditory and Reward ROI-to-ROI Connectivity

To separately evaluate the connectivity for auditory and reward networks at all four levels of liking and familiarity ratings, functional connectivity within the auditory network, within the reward network, and between auditory and reward networks were all significantly above chance, as shown in Figure 4A and 4C. Treating functional connectivity between auditory-auditory, reward-reward, and auditory-reward regions as dependent variables, a series of repeated-measures ANOVAs (three for liking and three for familiarity) were run to determine the main effect of time (pre-intervention vs post-intervention) rating and the time x rating interaction.

**Figure 4.**
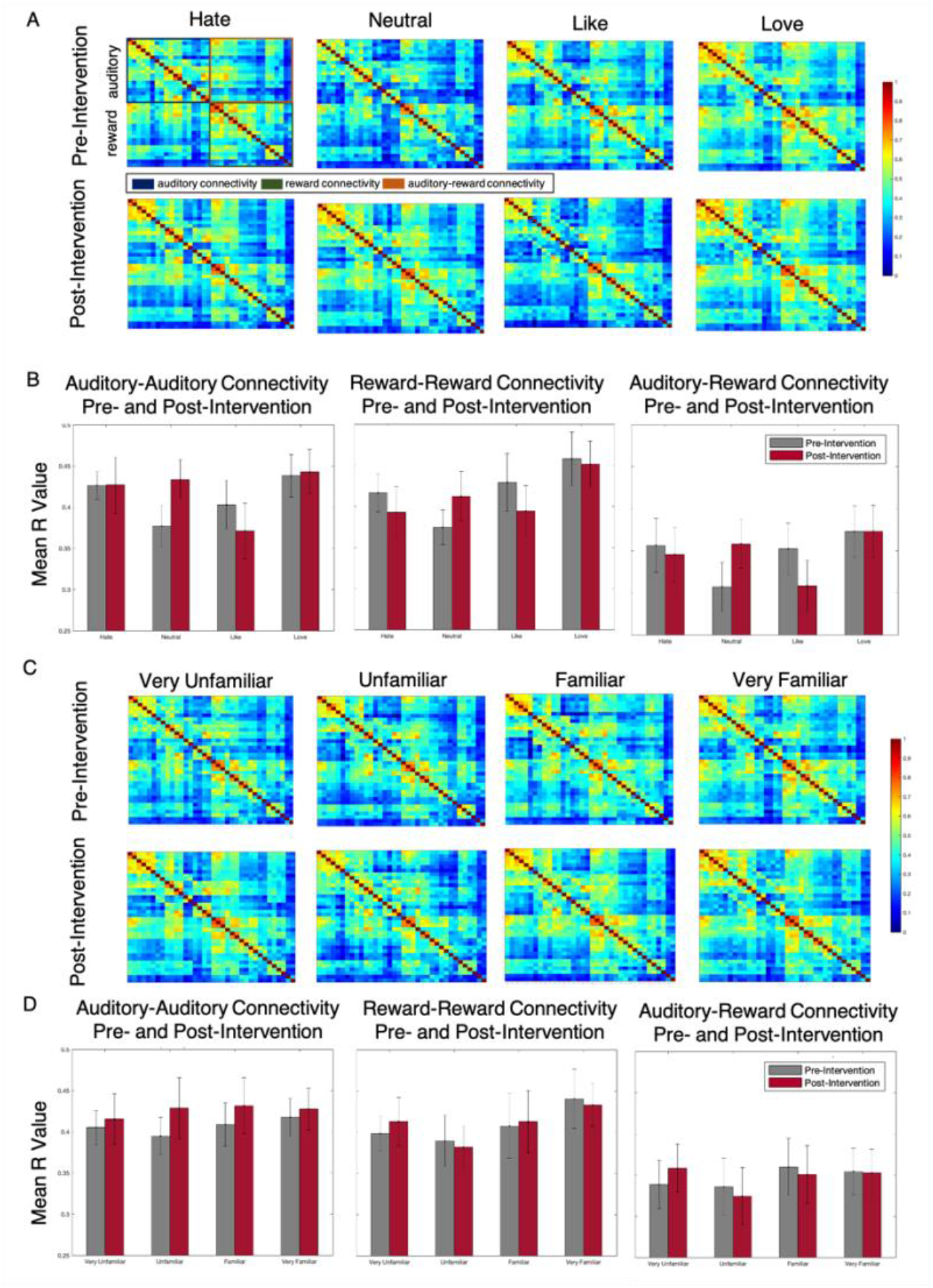
(A and C) Correlation matrices showing the relationship between auditory and reward regions for different liking (A) and familiarity (C) ratings. (B and D) Bar graphs depicting overall connectivity between auditory-auditory, reward-reward, and auditory-reward regions for different liking (B) and familiarity (D) ratings.

#### 3.5.1 Liking

For auditory-auditory connectivity, there were no significant main effects of time (F(1, 10) = 0.150, p = 0.707, 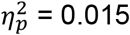) or ratings (F(3, 30) = 2.104, p = 0.121, 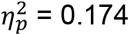). There was, however, a significant time x rating interaction (F(3, 30) = 3.068, p = 0.043, 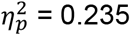), where hate and neutral ratings had higher beta-weights pre-intervention than post-intervention, but like and love had higher beta-weights post-intervention compared to pre-intervention. For reward-reward connectivity, there was an effect of rating (F(3, 30) = 2.923, p = 0.050, 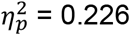). There was no significant main effect of time (F(1, 10) <0.001, p = 0.994, 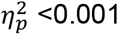) or time x rating interaction (F(3, 30) = 1.301, p = 0.292, 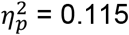). For auditory-reward connectivity, there was also no significant main effect of time (F(1, 10) = 0.001, p = 0.981, 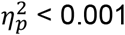) or rating (F(3, 30) = 1.647, p = 0.199, 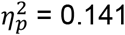) or time x rating interaction (F(3, 30) = 1.912, p = 0.149, 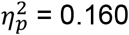). See Figure 4B.

#### 3.5.2 Familiarity

For auditory-auditory connectivity, there was no significant main effect of time (F(1, 8) = 0.264, p = 0.621 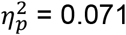) or rating (F(3, 24) = 0.608, p = 0.616, 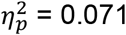) or time x rating interaction (F(3, 24) = 0.090, p = 0.965, 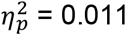). For reward-reward connectivity, there was no significant main effect of time (F(1, 8) = 0.282, p = 0.610, 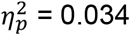) or rating (F(3, 24) = 2.476, p = 0.086, 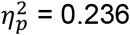) or time x rating interaction (F(3, 24) = 0.176, p = 0.912, 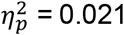). For auditory-reward connectivity, there was no significant main effect of time (F(1, 8) = 0.718, p = 0.421, 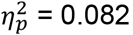) or rating (F(3, 24) = 0.923, p = 0.445, 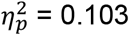) or time x rating interaction (F(3, 24) = 0.307, p = 0.820, 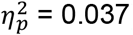). See Figure 4D.

### 3.6 Representational Similarity Analysis

A representational similarity analysis (methods outlined in Figure 5A) was conducted to compare how similarly the distinct ROIs and networks of interest (listed in Section 2.7.5) represented psychological characteristics of the musical stimuli, such as their novelty (well-known vs. novel stimuli), their selection (self-vs other-selected), and their liking and familiarity ratings, before and after intervention. Permutation testing comparisons of model RDMs to first-level fMRI-derived RDMs (Figure 5B) indicated a significant (p <0.05, T-max corrected) representation of novelty in the auditory network pre-intervention, but not post-intervention. In addition, there was significant representation of novelty in the left and right anterior and posterior STG, as well as representation of stimulus selection in the right posterior STG, and representation of liking in the anterior salience network at pre-intervention. Post-intervention, there were significant representations of stimulus selection in the left and right anterior and posterior STG and the basal ganglia, significant representations of novelty in bilateral posterior STG and left anterior STG, and significant representations of familiarity in both the basal ganglia and the language networks. Comparing pre-intervention and post-intervention fMRI-derived RDMs showed a significant effect of session in the representation of stimulus selection in the right executive control network (RECN), with a significant increase in relatedness post-intervention.

**Figure 5.**
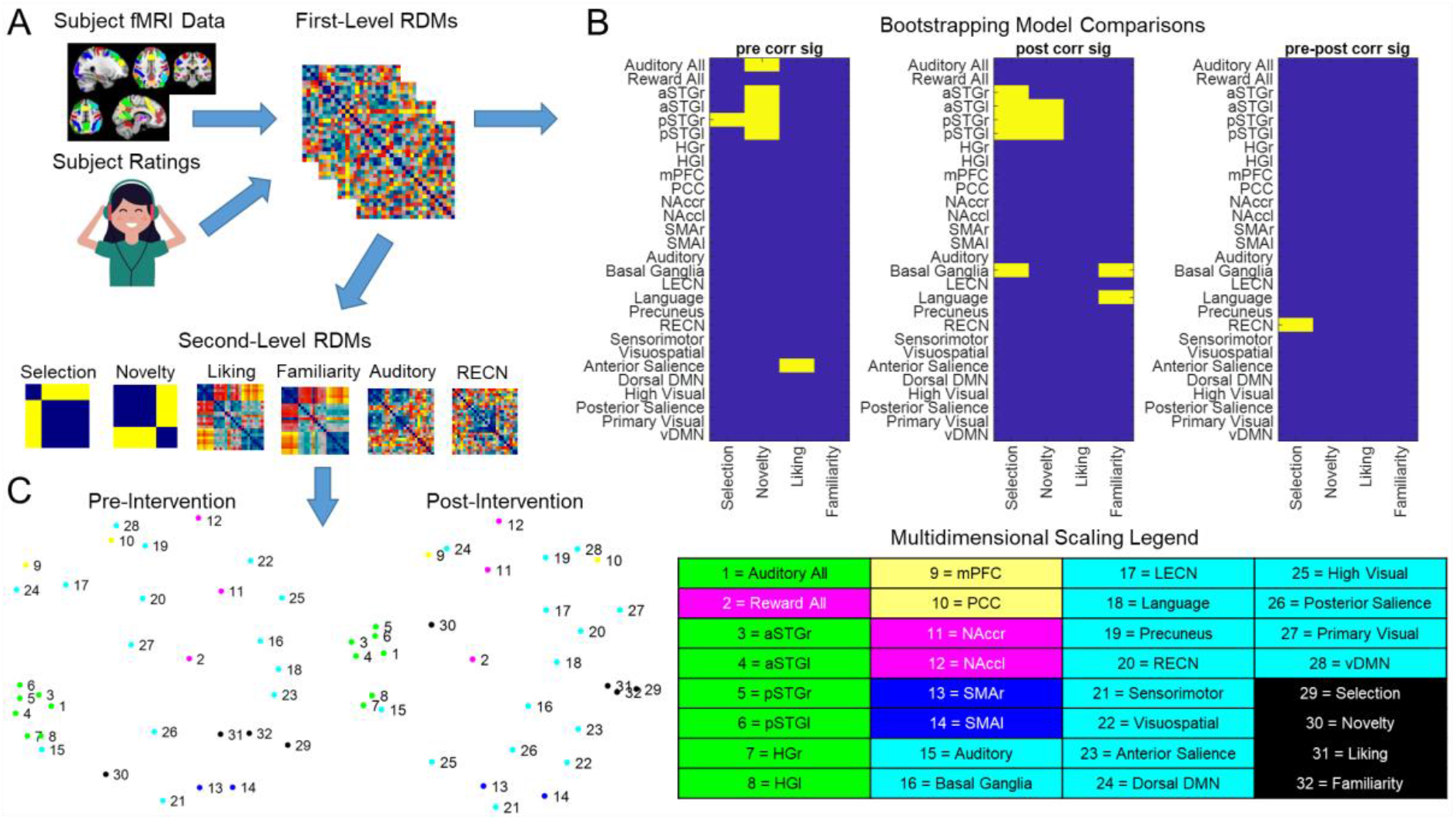
Representational Similarity Analysis. (A) Schematic of RSA processing pipeline. Participant-specific Ratings and task fMRI data are used to generate first-level representational dissimilarity matrices. First-Level RDMs are then used in 10,000-fold bootstrap resampling comparisons of models to functional RDMs for each participant. Resulting relatedness values are used in 50,000-fold t-max permutation testing (B) to determine significant (yellow) model-brain relationships at pre-intervention and post-intervention, as well as any significant effect of session (p<0.05). First-Level RDMs are then averaged across participants to generate second-level RDMs, which are used to produce second order multidimensional scaling visualizations of RDM relatedness (C).

Multidimensional scaling visualizations of second-level RDMs (Figure 5C) indicate a tight clustering of auditory (green) ROIs at both time points. These regions assort most closely with the model of novelty at both time points, and reward associated regions (pink) show a closer assortment with these regions post-intervention. A close relationship between models of liking, familiarity, and selection was also observed, and this relationship became even closer post-intervention. These models were most highly similar to bilateral SMA and the anterior salience network at pre-intervention, and they were most similar to anterior salience and language networks at post-intervention. Post-intervention representations of dorsal DMN and mPFC were also more closely related to bilateral nucleus accumbens as compared to pre-intervention. Overall, neural representations of musical stimuli were mostly similar before and after intervention, except that the reward network became more similar to the auditory network after intervention (Figure 5C), and the RECN showed increased differentiation between self-selected and other-selected stimuli, resulting in higher representational similarity with the stimulus selection model after intervention (Figure 5B, right panel).

## 4 Discussion

We present an fMRI study testing the effects of liking, familiarity, and self-selection on older adults. Furthermore, we present the first exploratory longitudinal fMRI results from an ongoing receptive music-based intervention on auditory-reward activity and connectivity in older adults. As predicted, musical excerpts that were self-selected, well-liked, and highly familiar were the most effective at driving the activity of the auditory and reward areas, especially overlapping in regions within the DMN. Functional connectivity of the auditory system and the mPFC increased over the course of the intervention and became more selective for self-selected and well-liked music. These results provide more support for the role of functional connectivity between the sensory and reward systems supporting preference, familiarity, and agency as shown by the effect of self-selection. We also see preliminary evidence for changes in brain connectivity following a music-based intervention; this has implications for the design and implementation of music-based interventions for special populations who have disrupted DMN or reward systems, including Alzheimer’s disease and depression, respectively.

Behavioral ratings of familiarity and liking, collected in the scanner before and after MBI, showed no overall difference between pre- and post-intervention liking ratings, but a stimulus-type by time interaction where ratings for self-selected stimuli were rated higher post-than pre-intervention, but well-known and novel stimuli were rated higher pre-intervention than post-intervention. Given that participants were listening to some of the same self-selected music as part of their intervention, this increase in liking could be a result of their increased familiarity due to repeated listening for a majority of the intervention. This similarity between familiarity and liking is broadly consistent with the fMRI data, which show many similar parametric effects between liking and familiarity in the auditory and DMN regions.

Univariate patterns of brain activity showed engagement of auditory and DMN regions. Both the parametric contrasts of liking and familiarity showed activity in auditory (STG, STS, MTG) and DMN (PCC) regions, with liking also engaging additional DMN regions (mPFC, TPJ/IPL) and familiarity also engaging a cluster in the precentral gyrus. While stimuli that were familiar and well-liked engage both the DMN and the auditory network, self-selected music was most effective at engaging additional DMN regions including the parahippocampal gyrus. This engagement of the DMN could be explained by the retrieval of autobiographical memories associated with the music that participants selected, as mPFC activity has been observed in music-evoked autobiographical memories [18]. These effects of liking and familiarity are largely consistent with previous reports, but this is the first study that parametrically tested different levels of liking and familiarity in the same study. In addition to treating liking and familiarity as continuous variables in a parametric analysis, we also separated them in our pairwise testing of auditory and reward connectivity, confirming significantly above-chance levels of functional connectivity within and between auditory and reward networks.

The effect of familiarity on mental representations of music has been shown on activity of the auditory network identified here [52-54], with effects on emotion [53] and mental representation of music [54]. While we did observe precentral gyrus engagement for familiarity, we did not observe significant activity in other auditory-motor regions, including the SMA. Precentral gyrus activity was only present for familiarity, but not for liking. This may suggest a motor engagement predictability that comes from the recognition of familiar musical elements, as has been shown from prior fMRI studies that involved listening to well-learned music [55]. Activity in auditory areas may reflect that the auditory system was able to form stronger predictions for familiar music than for unfamiliar music [52]. The DMN activity may reflect mind-wandering, or stimulus-independent thought [21, 56], including autobiographical memory and associations that may be elicited by music that is well-known to the listener.

Comparing the effects of liking against effects of familiarity, the TPJ/IPL and mPFC are two regions that were active for liking but not for familiarity. This distinction between liking and familiarity is important: the mPFC is part of both the DMN and the reward system, and it is consistently active during pleasurable experiences including but not limited to music [57]. Our results are consistent with Pereira et al. (2011), who also demonstrated frontal pole/mPFC activity for liked but not for familiar musical stimuli. The fact that it is active during the liking contrast and not the familiarity contrast, adds further support to mounting evidence that the mPFC has a unique role within the DMN, as a region that is common to the DMN and the dopaminergic system.

In addition to being part of the DMN [56], the TPJ and the mPFC have together been posited as a theory-of-mind network [58]. The TPJ/IPL is active during theory of mind tasks [59]. This role of TPJ/IPL in thinking about other minds fits well with the idea that musical experiences are fundamentally intertwined with the listener’s understanding of the intentions of the composer and the performer. It has been posited that “the best music” should have rich structure that can be inferred from its surface, and that the subjective value of music arises from an alliance between “compositional grammar” and the listener’s “listening grammar” [60]. While the relationship between the meaning of music and its subjective value is complex and abstract, the role of a theory-of-mind network may add to the claim that comprehensibility –– here, the understanding of other minds –– is necessary if not sufficient for subjective value in music.

Here, we see that while the TPJ/IPL and mPFC responded selectively to well-liked music, they showed even greater activity in the self-selected vs. other-selected contrast. The fact that self-selected music was more effective at engaging the theory-of-mind network, compared to music that was retroactively rated as well-liked, again points to a role of agency in determining the subjective value of music.

A strength of the present study is that we obtained fMRI data using a fast TR with many different kinds of musical stimuli, including some that were self-selected by the participant, and others selected by the researcher to span well-known musical selections as well as novel stimuli. The quality and quantity of data enabled more time-sensitive analyses of specific ROI data, including RSA to capture the relative distance between specific ROIs and functional networks, and between ROIs and experimentally-defined variables such as liking, familiarity, novelty, and self-selection. RSA confirmed that liking and familiarity for music are very similarly represented in the brain, and that both are strongly predicted by agency, or the effect of self-selection. Interestingly, the variables of liking, familiarity, and self-selection were more closely represented post-intervention than pre-intervention. Another important observation from RSA is that the reward network became more similar (closer in the MDS plot) to the auditory network after intervention. Novelty was represented differently from liking, familiarity, and self-selection. Relatedly, the RECN was the only network that showed a significant change in representation of stimulus features, specifically of the feature of selection. This is reflected in the increased dissimilarity in the RECN’s representation of the self-selected and novel stimuli post-intervention (Supplementary Materials Figure S6). In other words, the RECN was more representative of stimulus selection after intervention, in contrast to other networks such as the DMN. This pattern of increased dissimilarity between RECN and DMN that emerged after intervention converges with the well-replicated finding of anticorrelated activity between DMN and RECN at rest and during most cognitive tasks [61].

Seed-based connectivity showed an increase in functional connectivity from auditory regions to mPFC after intervention. This increase in connectivity is modulated by liking and familiarity, as well as by self-selection. This is borne out by between-participants correlations as well as within-participants beta time-series comparisons: Highest auditory-mPFC connectivity was observed across participants during liked and loved stimuli, especially after intervention. The beta time-series data showed an increasing differentiation by liking, familiarity, and self-selection after the intervention. In contrast, stimuli that were chosen to be well-known to the participants did not differ from novel stimuli in auditory-mPFC functional connectivity. Thus, the music-based intervention increased auditory-reward connectivity, but especially for some individuals and for some listening conditions. This is an important finding, as it is the first longitudinal demonstration of change in auditory-mPFC connectivity. Auditory-reward connectivity has been shown previously during the experience of pleasure when listening to music in task fMRI, PET, resting state fcMRI, and white matter connectivity as identified by DTI [39, 62, 63]. Auditory-reward connectivity has also been modulated pharmacologically [2] and with transcranial magnetic brain stimulation [64]. However, this is the first study to show a natural (i.e. as opposed to experimentally-induced) longitudinal change in auditory-reward connectivity after a receptive music-based intervention. By mindfully listening to music over the course of weeks, we have shown that participants can change their auditory-reward connectivity. As the reward system is important for many kinds of motivated behavior, these results have profound implications for the design of lifestyle interventions, such as music listening, which may affect motivated behavior through its underlying brain mechanisms.

The mPFC is a crucial hub across multiple networks in the brain, due to its membership in the dopaminergic reward network, in the DMN, and in coding for subjective value and a sense of ownership or self-referential processing [65, 66]. Rather than disentangling between these theoretical contributions to mPFC activity, the finding that music selectively engages mPFC may highlight the role that music has in all of these psychological functions. Our findings support the idea that music provides an auditory channel towards reward centers including the mPFC. This auditory channel towards the reward system is posited to be the evolutionary basis for the role of music for social bonding [15].

From a clinical standpoint, the mPFC and PCC are most sensitive to changes in functional connectivity in Alzheimer’s disease [21]. mPFC activity and connectivity are also disrupted in psychiatric diseases, including depression and schizophrenia [22]. Thus, these results suggest that MBI may help those with depression and AD, and possibly those with neurodegenerative and psychiatric diseases more generally, by mitigating aberrant network changes through modifying activity and connectivity of the mPFC. While these results suggest that MBI is effective in changing auditory-reward connectivity, one caveat is that the present longitudinal study is not a randomized controlled trial; thus the results could be explained by time-dependent changes that are extraneous to music listening. However, given that the older adult population sampled here is unlikely to have spontaneously increasing brain connectivity in these areas without a focused intervention, and given that auditory-reward connectivity is a relatively well-established set of regional hypotheses from previous research [5, 46, 57], we expect that the observed changes here point to the longitudinal effects of music listening. Nevertheless, the next phase of this study will include a auditory but non-musical control intervention to help isolate the roles of specific musical features on changes in auditory-reward activity and connectivity.

This study aims to provide a preliminary look at the effects of an MBI on older adults. Though the sample size for this study (N=16) is relatively small at this point, these results offer a promising direction for research on MBIs and their effect on the aging brain. The COVID-19 pandemic significantly impacted recruitment for this study, limiting trajectory of our recruitment goals and the participants who felt comfortable coming into the lab. Future directions of this study will continue to build on the promising results reported here.

More generally, the findings of this study suggest that participants should choose the music to be applied in a given intervention or music therapy treatment program, in order to leverage the effects of agency on engagement of brain functions. Allowing individuals to self-select music will also necessitate collaborations between neuroscientists and music therapists to implement MBIs in ways that respect the cultural dependence and personal salience of musical experiences [67]. By demonstrating the ability of music to engage the connection between auditory and reward systems in healthy older adults, our findings shed light on potential therapeutic effects of music listening on healthy aging.

## 5 Acknowledgments

Supported by the Grammy Foundation, the Kim and Glen Campbell Foundation, NSF-STTR #2014870, NSF-CAREER award #1945436, and NIH R21AG075232 to PL. We thank all our participants, NUBIC, and Juliet Davidow and the Northeastern University MRI Users Group for helpful comments on a previous version of this manuscript.

